# Circadian coupling of mitochondria in a deep-diving mammal

**DOI:** 10.1101/2023.11.06.565796

**Authors:** Chiara Ciccone, Fayiri Kante, Lars P. Folkow, David G. Hazlerigg, Alexander C. West, Shona H. Wood

## Abstract

Regulation of mitochondrial oxidative phosphorylation is essential to match energy supply to changing cellular energy demands, and to cope with periods of hypoxia. Recent work implicates the circadian molecular clock in control of mitochondrial function and hypoxia sensing. Since diving mammals experience intermittent episodes of severe hypoxia, with diel patterning in dive depth and duration, it is interesting to consider circadian - mitochondrial interaction in this group. Here we demonstrate that the hooded seal (*Cystophora cristata*), a deep diving Arctic pinniped, shows strong daily patterning of diving behaviour in the wild. Cultures of hooded seal skin fibroblasts exhibit robust circadian oscillation of the core clock genes *per2* and *arntl*. In liver tissue collected from captive hooded seals, expression of *arntl* was some 4-fold higher in the middle of night than in the middle of the day. To explore the clock-mitochondria relationship, we measured the mitochondrial oxygen consumption in synchronized hooded seal skin fibroblasts and found a circadian variation in mitochondrial activity, with higher coupling efficiency of complex I coinciding with the trough of *arntl* expression. These results open the way for further studies of circadian - hypoxia interactions in pinnipeds during diving.

**SUMMARY STATEMENT:** A functional clockwork and circadian variation in mitochondrial complex I efficiency is demonstrated in skin fibroblasts from the deep diving hooded seal.

## INTRODUCTION

The ability of animals to predict and prepare for daily changes in environmental demands relies on the presence of an intrinsic physiological system: the circadian clock. The molecular basis of the circadian clock in mammals is a transcription-translation negative feedback loop (TTFL). The positive limb of this loop depends on the dimerization of the transcriptional activators CLOCK and ARNTL (also known as BMAL1), while the negative limb depends on the formation of a transcriptionally repressive complex containing PER and CRY (Buhr and Takahashi, 2013). Circadian dynamics emerge through the autoregulatory effects of PER and CRY: the CLOCK/ARNTL complex activates transcription of PER and CRY which, after dimerization, translocate into the nucleus and repress their own transcription. This creates a loop with a period of around 24-hours and therefore called *circadian*.

One of the roles of the circadian clock is to coordinate the metabolic transcription network, presumably in order to optimize mitochondrial metabolism to the daily changes (Neufeld-Cohen et al., 2016; Turek et al., 2005). In mice, mitochondrial oxidative phosphorylation (OXPHOS) oscillates on a daily manner and in concomitance with the expression of some rate-limiting mitochondrial enzymes (Neufeld-Cohen et al., 2016). OXPHOS results from the activity of enzymatic complexes (I-IV) located on the inner mitochondrial membrane (IMM). These complexes are responsible for the transport of electrons through the IMM and are therefore referred to as the Electron Transport System (ETS). Complexes I and II catalyse the transfer of electrons from the two tricarboxylic acid (TCA) cycle products, NADH and FADH_2_, to complex III via ubiquinone. The electrons are next transferred via cytochrome c to complex IV where oxygen (O2) acts as the final electron acceptor. The electron transfer process enables complexes I, III and IV to pump protons (H^+^) from the mitochondrial matrix into the intermembrane space (Hosler et al., 2006), thus creating the electrochemical gradient that drives ATP production by ATP synthase, also referred to as complex V (Chance and Williams, 1956). Variations in the activity of these complexes generate variations in mitochondrial OXPHOS and ATP production.

In mice with a dysfunctional circadian clock, the daily cycle of OXPHOS is abolished, suggesting that there is an intrinsic dependence of the mitochondrial respiration complexes on the circadian clockwork (Neufeld-Cohen et al., 2016). Deletion of *arntl* in C2C12 myotubes causes a reduction of both OXPHOS and extracellular acidification rate (an indicator of glycolytic rate), but deletion in embryonic stem cells increases OXPHOS while reducing glycolytic rate (Ameneiro et al., 2020; Peek et al., 2017). This suggests there is tissue specificity in the effect of “circadian clock knockout” on mitochondrial responses.

O_2_, the final electron acceptor of the ETS, constitutes a fundamental factor in mitochondrial regulation. In mouse and human cell cultures, hypoxia (O_2_ deficiency) leads to a switch from OXPHOS-based metabolism to anaerobic glycolysis (Kim et al., 2006; Semenza et al., 1994; Wang and Semenza, 1993). In mice, this hypoxia-driven mitochondrial switch to enhanced glycolytic metabolism involves hypoxia inducible factor 1α (HIF-1α), and its interaction with the clock gene, ARNTL (Peek et al., 2017). In addition, several studies have highlighted a circadian clock – mitochondria and hypoxia interaction (Adamovich et al., 2017; Manella et al., 2020; Peek et al., 2017), however, such links have only been demonstrated in rodent species that do not normally experience severe hypoxia. It is therefore of interest to investigate whether circadian – mitochondrial interactions are also present in a diving mammal, which through its behaviour frequently experiences episodes of hypoxia.

Diving mammals have evolved numerous hypoxia coping strategies including enhanced O_2_ storage and O_2_ carrying capacity with blood volume and hemoglobin and myoglobin levels being higher than in other species (e.g., Blix, 2018; Davis, 2014). These adaptations combine with strict O_2_ economy by way of cardiovascular adjustments (bradycardia and peripheral vasoconstriction) and hypometabolism (Blix et al., 1983; Burns et al., 2007; Lenfant et al., 1970; Scholander, 1940). For the majority of dives this means that animals are within their aerobic dive limit (ADL). Nevertheless, some deep-diving seal species (Weddell seals (*Leptonychotes weddellii*), northern elephant seals (*Mirounga angustirostris*), hooded seals *(Cystophora cristata*)) appear to repeatedly perform dives that exceed their calculated ADL (Folkow et al., 2008; Meir et al., 2009; Qvist et al., 1986). Arterial partial O_2_ pressures (tension) of 10-20 mmHg have been recorded in freely diving seals (Meir et al., 2009; Qvist et al., 1986), these values are far below what is considered as a critically low arterial partial O_2_ pressure for adequate brain function (25-40 mmHg) (Erecińska and Silver, 2001). Therefore, deep-diving species are likely to experience severe hypoxia on a regular basis.

There are several reports of seasonal and daily variations in diving behaviour of different pinnipeds species (Bennett et al., 2001; Folkow and Blix, 1999; Nordøy et al., 2008; Photopoulou et al., 2020). Among them, the hooded seal is known for its remarkable diving capacity (Folkow and Blix, 1999): dives exceeding durations of 1 hour and depths of 1600 meters have been recorded (Andersen et al., 2013; Vacquie-Garcia et al., 2017), making this species a good research model for hypoxia tolerance and oxidative stress in mammals (e.g., Fabrizius et al., 2016; Folkow et al., 2008; Geiseler et al., 2016; Geßner et al., 2020; Hoff et al., 2016; Hoff et al., 2017; Mitz et al., 2009; Vázquez-Medina et al., 2011). The hooded seal dive duration and depth are reportedly greater during the day than at night-time (Andersen et al., 2013; Folkow and Blix, 1999), suggesting that there may be a daily rhythm in exposure to severe hypoxia. We hypothesise that, in addition to other diving adaptations, a diving mammal may use circadian - mitochondrial interactions to tolerate variations in oxygen availability. However, the circadian molecular clock is yet uncharacterised in pinnipeds, and mitochondrial activity has not been measured in the deep diving hooded seal.

Here, we provide the first molecular characterisation of the circadian clock in the hooded seal and explore the clock-mitochondria relationship, showing that mitochondrial activity relates to circadian clock phase. These results open the way for further studies of circadian and (diving-induced) hypoxia interactions in pinnipeds.

## MATERIALS AND METHODS

### Diving behaviour analysis

Data collected by the Norwegian Polar institute in 2007 and 2008 as part of the MEOP (Marine Mammals Exploring the Ocean Pole to Pole) programme were provided by Dr. M. Biuw and used for diving behaviour analysis based on dive data for 12 adult and 8 juvenile hooded seals of both sexes (Vacquie-Garcia et al., 2017, <u>10.21334/npolar.2017.881dbd20</u>, file name: dive_2007.txt, dive_2008.txt). Data collection was funded by the Norwegian Research Council (grant number 176477/S30) and the Norwegian Polar Institute. Conductivity-Temperature-Depth Satellite relay Data Loggers (CTD-SRDLs) (Sea Mammal Research Unit, University of St Andrews) were glued to the fur on the back of the neck of the seals. Data were collected and transmitted for every 6-hour period and included the following variables: dive duration, time of dive end, maximum depth, location (latitude and longitude), and time spent at surface after the dive. Knowing dive duration, time of dive end and time spent at surface, it was possible to infer the starting time for every single dive. Starting time was taken to represent the start of a ‘diving event’, which then formed the basis for analyses of diurnal changes in dive duration and depth.

Hooded seal pups perform shallower and shorter dives than adults (Folkow et al., 2010). Our analyses therefore focused on the 12 adults (3 males and 9 females) that were represented in the data set.

After having evaluated the dive duration distribution, we decided to analyse only dives between 2 minutes and the 95% percentile of the maximum dive duration of each individual. This allowed us to exclude outliers from further analysis.

All the following analysis were done in RStudio (4.2.1 version). To verify the presence of a daily oscillation in the hooded seal diving behaviour, a cosine curve with a 24-h period was fitted to the data through the function *cosinor.lm* in the package *cosinor*. To test whether the cosine model significantly represented the data, we used the *cosinor.detect* function in the package *cosinor2*. Calculations were repeated for each seal and for each month separately. The function *cosinor* in the package *card* was used to define the Midline Statistic of Rhythm (MESOR): this is a circadian rhythm-adjusted mean which gives an estimate of the average value of an oscillating variable. The MESOR was used to identify different trends in diving duration throughout the year. Using the information about latitude and longitude, it was possible to calculate time of sunrise and sunset for each day of every month through the *sunriset* function in the *maptools* package.

To compare dive durations at different times of day we binned the data by hour and calculated the hourly mean dive duration.

### Animals used for tissue sampling

Hooded seals (*Cystophora cristata*) were captured in their breeding colonies on the pack ice of the Greenland Sea, at ∼71°N and ∼019° W, during a research/teaching cruise with the R/V Helmer Hanssen in late March 2018, under permits of relevant Norwegian and Greenland authorities. Six live seals were brought to the Department of Arctic and Marine Biology (AMB) at UiT – The Arctic University of Tromsø, Norway, where they were maintained in a certified research animal facility (approval no. 089 by the Norwegian Food Safety Authority (NFSA)). Seals were euthanized (in 2019, as juveniles, at age ∼10 mo.) for purposes other than the present study, in accordance with a permit issued by NFSA (permit no. 12268): the seals were sedated by intramuscular injection of zolazepam/tiletamine (Zoletil Forte Vet., Virbac S.A., France; 1.5 – 2.0 mg per kg of body mass), then anaesthetized using an endotracheal tube to ventilate lungs with 2 – 3 % isoflurane (Forene, Abbott, Germany) in air and, when fully anaesthetized, they were euthanized by exsanguination via the carotid arteries.

During another research cruise in March 2019, additionally 6 weaned pups were captured (NFSA; permit no. 19305) and brought to AMB, UiT. Before culling and tissue sampling (February 2020, at age 11-12 mo.), the animals were exposed to 18 days of 12 h light and 12 h darkness (12L;12D). In this experiment, we defined Zeitgeber Time 0 (ZT0) as the time when lights went on. Euthanasia was performed as explained above at ZT6 (light phase) for three seals and at ZT18 (dark phase) for the other three.

### Tissue sampling

Skin biopsies were collected postmortem from the seal captured in March 2018 between the digits of the hind flipper with a biopsy punch (6 mm, 33-36-10, Miltex Inc., York) and processed for culturing of fibroblasts as described below. Samples from kidneys and liver were collected from the 6 seals held under 12L;12D light conditions. Tissues were minced with a sterile scalpel blade and placed in 4 ml of RNAlater (AM7021, Thermo Fisher) in a 1:10 ratio. Samples were stored at 4°C for 24 h and then moved to -20°C until mRNA extraction.

### Culture of hooded seal skin fibroblasts

Skin biopsies were processed as described elsewhere (Du and Brown, 2021). Briefly, biopsies were placed in collection medium (Dulbecco’s Modified Eagle Medium (DMEM, D5796, Sigma) + 50 % Fetal Bovine Serum (FBS, F7524, Sigma) + 1% penicillin-streptomycin (Pen-Strep, P4458, Sigma)) and then moved into 6-well-plates containing digestion medium (DMEM + 10% FBS + 1% amphotericin B (A2942, Sigma)) and 10% liberase (Roche, 05 401 119 001) for 8-9 hours at 37° C/5% CO_2_. Tissue biopsies and digestion medium were then transferred to a tube containing warm Dulbecco’s Phosphate Buffered Saline (PBS, D8537, Sigma), plates were rinsed again with PBS to collect all the tissue fragments. Samples were centrifuged at 200 g for 5 minutes. The pellet was resuspended in culture medium (DMEM + 20% FBS + 0.1% gentamycin (15710, Invitrogen)) and placed in fresh 6-well-plates. Fragments were overlaid with a Millicell Cell Culture insert (Cat. PICMORG50 Millipore): 1.5 ml of culture medium was added to the interior of the insert and 0.5 ml to the exterior. Plates were incubated for about 2 weeks at 37° C/5% CO_2_. Medium was changed every 3-4 days.

When cells covered approximately 80% of the plates surface (80% confluency), they were trypsinised (Trypsin-EDTA solution, T4049, Sigma), resuspended in culture media and centrifuged at 400 g for 5 minutes. Pelleted cells were then suspended in 3 ml of freezing medium per T75 culture flask (culture medium + 10% Dimethyl sulfoxide (DMSO, D5879, Sigma)) and progressively cooled in a cryocooler overnight at - 80° C. Cells were then moved to liquid nitrogen for long-term storage.

A few days before the start of the experimental procedures (temperature cycling and respirometry) cells were thawed at 37°C for 1-2 minutes and then added to 10 ml of pre-warmed medium (DMEM + 20% FBS + 1% Pen-Strep). The solution was centrifuged at 300 g for 5 minutes. The pellet was resuspended in 12 ml of medium and transferred to a T75 culture flask. When cells reached 80 % confluency, they were trypsinised and re-plated into three different plates (1 T175, 1 T75 and 1 6-well-plate) for each sampling timepoint.

### Temperature cycling treatment

According to established protocols, temperature cycling was used to synchronise the circadian clock of our primary fibroblast cell cultures (Brown et al., 2002; Buhr et al., 2010). Cells were exposed to five consecutive 24h-cycles of temperature alternations between 12-h at 36.5°C and 12-h at 39.5°C. In all the temperature cycling experiments, we defined ZT0 as the time of the transition from 36.5 to 39.5°C. Cells were collected from the sixth cycle with 4-hour intervals over 2 cycles (Fig. S1A). For mRNA, 6-well plates were washed with PBS and directly frozen on dry ice before storage at - 80°C. For the mitochondrial O_2_ consumption measurements, cells grown in T175 flasks were sampled with an 8-hour interval, thus giving data around 1 full 24h-cycle with only one repetition for each sampling timepoint (Fig. S1A). The presence of phenol red in the culture medium was used as standard pH indicator in the cultures: if changes in pH were detected, the medium was changed at the end of the third cycle.

In a second experiment, cells were kept under temperature cycle for 3 cycles. Cells were sampled over a 48-hour period, at 36.5°C for the first 12 hours and at 39.5°C for the remaining 36 hours (Fig. S1B).

### mRNA extraction, cDNA conversion

For both tissues and cells, RNA was extracted using RNeasy mini kit (74104, Qiagen) following the manufacturer’s instructions. Tissue samples (∼100mg) were thawed then inserted in a low bind tube containing 600 µl RLT buffer + 6 µl β-mercaptoethanol with a metal bead. The tube was shaken in a Tissue Lyser for 6 minutes at 20 shakes/second. Cell plates were taken out of the - 80°C freezer and the contents of each well was collected in 350 µl of RLT buffer before being transferred to a QIAshredder column (79654, Qiagen). The column was centrifugated for 2 minutes at full speed and 350 µl of 70% ethanol were added to the lysate. Lysates were treated with DNase I (79254, Qiagen) and heated to 37°C for 10 minutes. Enzymatic reaction was stopped by adding EDTA at a final concentration of 0.05 M in each tube at 75°C for 10 minutes. Samples were then treated with cDNA buffer and enzyme reverse transcriptase, heated for 1 hour at 37°C and cooled down to 4°C. The cDNA samples were then stored at - 20°C until further use.

### qPCR

Seal gene sequences for *arntl*, *per2* and housekeeping genes *tbp* and *ppib* were identified by aligning in BLAST dog (*Canis lupus*) genes sequences against the Weddell seal (*Leptonychotes weddellii*) genome (https://www.ncbi.nlm.nih.gov/assembly/GCF_000349705.1/). Primer specificity was confirmed by cloning PCR amplicons into Zero Blunt TOPO vector and sanger dideoxy sequencing (BigDye). Only primers with efficiency above 90% and a single product were used (Table 1). Reagents from the Promega GoTaq qPCR Master Mix kit (A6001, Promega) were used to perform qPCR. The Bio-Rad manager software was used to control the Bio-Rad CFX Connect Real_Time PCR system. The qPCR template used started with 2 minutes at 95°C. It then repeated 39 temperature cycles, each of: 15 seconds at 95°C, 15 seconds at 57°C and 1 minute at 60°C, where fluorescence was measured. It finished with 5 seconds at 65°C and then a final constant temperature of 95°C. The cycle threshold (CT) values of each gene were analysed using the ΔΔCT (2^–ΔΔCt^) method (Livak and Schmittgen, 2001) against housekeeping genes *ppib* and *tbp*.

**Table 1.**
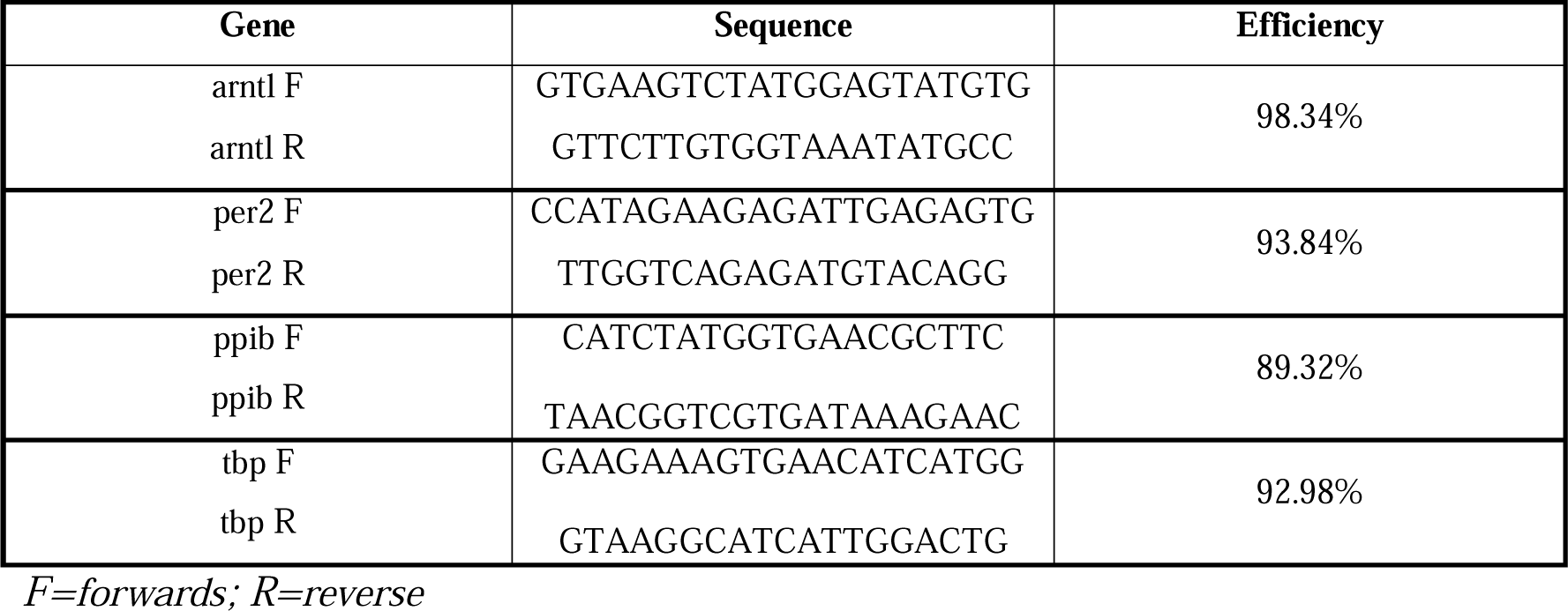
Primers sequences used for qPCR: efficiency calculated at 57° C.

### Mitochondrial oxygen consumption measurements

Mitochondrial oxygen consumption was measured with two O2k high-resolution respirometers (Oroboros Instruments, Innsbruck, Austria) and data were recorded in real-time using the Oroboros DatLab software (Oroboros Instruments).

Due to the duration of the respiration measurement protocol, and limited availability of Oroboros respirometers, it was possible to process samples only six times over 48 hours. After exposure to temperature cycling treatment, cells were collected by trypsinisation, resuspended in culture medium, and centrifuged at 300 g for 5 minutes. Pellet was then resuspended in the respiration medium MiR05 (0.5 mM EGTA, 3 mM MgCl_2_, 60 mM K-lactobionate, 20 mM taurine, 10 mM KH_2_PO_4_, 20 mM HEPES, 110 mM sucrose, 1g/L BSA; pH = 7). Cells were added to the Oxygraph chambers at a concentration of 0.6x10^6^ cells/ml with a final volume of 2.1 ml in each chamber.

After measurement of basal respiration (Routine), cells were permeabilised with digitonin (3 µg/ml). Different mitochondrial states were assessed by injecting the following chemicals in the chambers: pyruvate, malate, ADP, glutamate, succinate, cytochrome c, rotenone, oligomycin and carbonyl cyanide p-trifluoro-methoxyphenyl hydrazone (FCCP). Optimal concentration of both digitonin (Doerrier et al., 2018) and FCCP were determined in pilot experiments (Fig. S2). Table 2 gives an overview of the chemicals injected and the respiratory states measured. The initial O_2_ concentration in the chamber was ∼200 µM. Along the experiment, the Oxygen concentration decreased, down to ∼80-100 µM. Figure S3 shows an original oxygraph obtained during the mitochondrial respiration measurements.

**Table 2.** List of chemicals used during the oxygen consumption measurements and relative mitochondrial state.

| Chemical | Cat. No. and Supplier | Final Concentration | State | Description |
| --- | --- | --- | --- | --- |
| - | - | - | Routine | Basal respiration without addition of any substrate. |
| Digitonin | D5628, Sigma | 3 µg/ml | - | Cell permeabilization |
| Pyruvate | P2256, Sigma | 5 mM | LeakN | Leak state at Complex I in the absence of ADP (inactive ATP synthase): non-phosphorylating state where protons leaking through complex I are not used for ATP production. |
| Malate | M1000, Sigma | 2 mM |  |  |
| ADP | 117105, Calbiochem | 2.5 mM | OXPHOS (PM – P) | Oxidative phosphorylation through complex I with pyruvate and malate as substrates |
| Glutamate | G1626, Sigma | 10 mM | OXPHOS (CI – P) | Oxidative phosphorylation through complex I with pyruvate, malate and glutamate as substrates |
| Succinate | S2378, Sigma | 10 mM | OXPHOS (CI + CII – P) | Oxidative phosphorylation through complex I and complex II with pyruvate, malate, glutamate, and succinate as substrates |
| Cytochrome c | C7752, Sigma | 10 µM | - | Mitochondrial integrity test |
| Rotenone | R8875, Sigma | 0.5 µM | OXPHOS (CII – P) | Oxidative phosphorylation through complex II after inhibition of complex I |
| Oligomycin | O4876, Sigma | 10 nM | LeakOmy | Leak state at complex II after inhibition of ATP synthase: non phosphorylating state |
| FCCP | C2920, Sigma | 1 µM | Uncoupled state (CII – E) | Uncoupled state measuring maximal mitochondrial capacity |

Each experiment was done with four replicate respiratory chambers, except for ZT13 and ZT21, which have three replicates. The fourth replicate was excluded because the cells did not permeabilize, therefore the mitochondria did not respond to the treatments.

The data obtained for each oxygraphic chamber were first normalised to the specific number of cells within that chamber (expressed as multiples of 10^6^ cells) in the DatLab software (Fig. S4), and then normalised also to the Routine respiration of that same chamber; this was done to eliminate any sample-specific variation. Finally, measurements at different timepoints for each mitochondrial state were normalised to cycle mean, to reveal any rhythmicity. The cycle mean was calculated as the average value for each mitochondrial state; single measurements were then divided by the cycle mean, to be represented by ratios as oscillations around 1 (Fig. S5). A significant difference between timepoints was found for all the mitochondrial states, both before and after normalisation to cycle mean (one-way ANOVA analysis).

The *leak state* is defined as the mitochondrial respiration in the presence of fuel substrates and in the absence of ADP. It reflects proton leak across the inner membrane that does not result in ATP production (Chance and Williams, 1956; Gnaiger, 2009; Perry et al., 2013). It can be used to determine the coupling efficiency of the different ETS enzymes through the respiratory control ratio (RCR), measured as OXPHOS/LEAK (P/L), where OXPHOS corresponds to mitochondrial respiration in the presence of substrates and saturating ADP (Chance and Williams, 1956; Gnaiger, 2009). In the literature, the RCR is preferably expressed as OXPHOS coupling efficiency (1 – RCR^-1^), with values between 0 and 1, where 1 corresponds to fully coupled mitochondria (Doerrier et al., 2018; Gnaiger, 2020). Accordingly, we calculated complex I coupling efficiency using this formula:

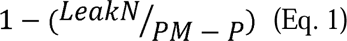

Where LeakN corresponds to *leak state* at complex I and PM-P to complex I OXPHOS, both with pyruvate and malate as substrates (Table 2).

### Lentiviral vector construction

Circadian clock gene activity was recorded using the vectors pLV6-Bmal-luc (Addgene plasmid #68833) and pLV6-Per2-luc. The latter consisted of the pLV6 backbone and a mouse *per2* promoter with adjacent luciferase sequence contained in the pGL3 basic E2 vector (Addgene plasmid 48747). To ligate the Per2:luciferase reporter with pLV6 backbone we designed a restriction cloning approach shown to be efficient in large plasmids using the QuickChange Lightning Site-Directed Mutagenesis (SDM) kit (Agilant, 210518) according to manufacturer’s instructions (Zhang and Tandon, 2012). Briefly, the SDM kit was used to create an extra BamHI cutting site 5’ on the pLV6-Bmal-luc vector. Mutated plasmid was transformed with stable competent *E. coli* (NEB, C3040H) according to the manufacture’s manual. Cells were grown on LB agar plates (40g/l LB-agar (Sigma-Aldrich, L3147), 100 µg/ml Ampicillin (Sigma-Aldrich, A9393)) and single clones were cultivated in LB broth overnight (25g LB (Sigma-Aldrich, L3522), 100 µg/ml Ampicillin). We used a Qiagen miniprep kit (Qiagen, 27104) to extract plasmid DNA and the mutation was confirmed through whole-plasmid sequencing (Plasmidsaurus, https://www.plasmidsaurus.com/). After the mutation was detected, the pLV6-Bmal-luc vector was digested with BamHI (NEB, R3136S) then treated with quick CIP (NEB, M0525S) to avoid self-ligation. The digest was run on an 1% Agarose gel (TAE buffer, 0.25 µg/ml EtBr) and the the QIAquick Gel extraction kit (Qiagen, 28704) was used to extract the pLV6 backbone according to the manufacturer’s instructions. Since the Per2 promoter and luciferase in the pGL3-Per2-luc vector is already flanked by BamHI cutting sites, the pGL3-Per2-luc vector was directly digested with BamHI for Per2:luciferase sequence extraction.

Finally, the pLV6 backbone and the Per2:luciferase fragment were ligated with T4 DNA ligase (NEB, M0202S) according to the manufacture’s manual. Vectors extracted from transformed single colonies were sequenced by Plasmidsaurus and checked for correct ligation and sequence. The resulting vector is referred to as pLV6-Per2-luc.

Lentiviral vectors were constructed according to (Du and Brown, 2021). Briefly, HEK293T cells were incubated with pMD2G (#12259, Addgene), psPAX2 (#12260, Addgene), and the two transfer vectors. Supernatant was collected 24 and 48 hours after incubation and transfection and filtered through a 0.2 µm filter. Filtered supernatant was aliquoted and frozen at -80°C until use.

### Generation of a stable cell line expressing lentiviral vector and bioluminescence recordings

Two ml of medium containing lentivirus were added to a T25 flask containing hooded seal fibroblasts at 50% confluency, with 3 ml of DMEM + 20% FBS. Cells were incubated overnight and then the media was changed to clean cells from the virus. After 3 days, medium was changed to DMEM + 20% FBS + 10 µg/ml blasticidin (R21001, Gibco) selection antibiotic (cells were passaged in a T75 flask at this point). From the T75 flask, 10 3.5-cm-wells were seeded and cultured in DMEM + 20% FBS + 10 µg/ml blasticidin. One week before starting the temperature cycling, medium was changed to recording medium (DMEM without phenol red + 0.47% NaCO_3_ + 1% HEPES buffer + 0.25 % Pen-Strep + 5% FBS + 10 µg/ml blasticidin + 0.1 mM luciferin) (Yamazaki and Takahashi, 2005). Prior to the experiment, medium was again changed, and wells were sealed with parafilm and placed in a Photon multiplier tube (Lumicycle, Hamamatsu) for recording.

On a separate experiment, after a 2-days incubation at 37° C/5% CO_2_ in recording medium, transfected cells were synchronized with dexamethasone (100 nM, D4902, Sigma) for 30 min. Thereafter, cells were washed twice with PBS and medium was changed to normal recording medium. Cells were then sealed and placed in the Photon multiplier tube for recording.

### Statistical analysis

All statistical analyses were performed in RStudio using one-way ANOVA and post comparison Tukey HSD test. Statistical analysis of coupling efficiencies was also performed using the non-parametric test Kruskal-Wallis across all the ZTs and Mann-Whitney U-test for peak and through comparison. The analysis of circadian oscillations for clock genes mRNA expression was done using JTK cycle, a non-parametric algorithm designed to detect cyclic patterns in datasets with regular intervals between measures (Hughes et al., 2010). All data are represented as mean ± standard error (SEM). Periods of the photon multiplier tube recordings were analysed by fitting a damped sine wave in GraphPad Prism 8 (version 8.0.2). Principal Component Analysis (PCA) of ZTs was performed across the 7 measured mitochondrial states using the *prcomp* function in RStudio. Ellipses were drawn with a 0.80 confidence interval.

## RESULTS

### Seasonal and daily variations in hooded seal diving behaviour

We used the data collected between 2007 and 2008 by the Norwegian Polar Institute (see Vacquie-Garcia et al., 2017) to perform a high time resolution analysis of hooded seal diving behaviour. Initial analysis revealed that the maximum dive duration was 87.25 minutes for males (at ∼09:00 in January 2007, adult male Mj98) and 53.25 minutes for females (at ∼10:00 in April 2008, adult female F175). For most seals, cosinor analysis of diving duration data showed that there is a significant relationship between dive duration and time of the day with a 24-h period throughout the year (Table 3). Figure 1 shows this relationship for a male adult hooded seal (ID: M171, August 2007 – April 2008). While dives were observed in the night, the average dive duration was consistently higher during daytime (05:00 – 16:00) for most of the months, and very few night-time dives were longer than 1 hour (Fig. 2). Therefore, by investigating the hourly distribution of diving durations, we were able to confirm the presence of a diurnal pattern in the hooded seal diving behaviour (Fig. 1, 2).

**Figure 1.**
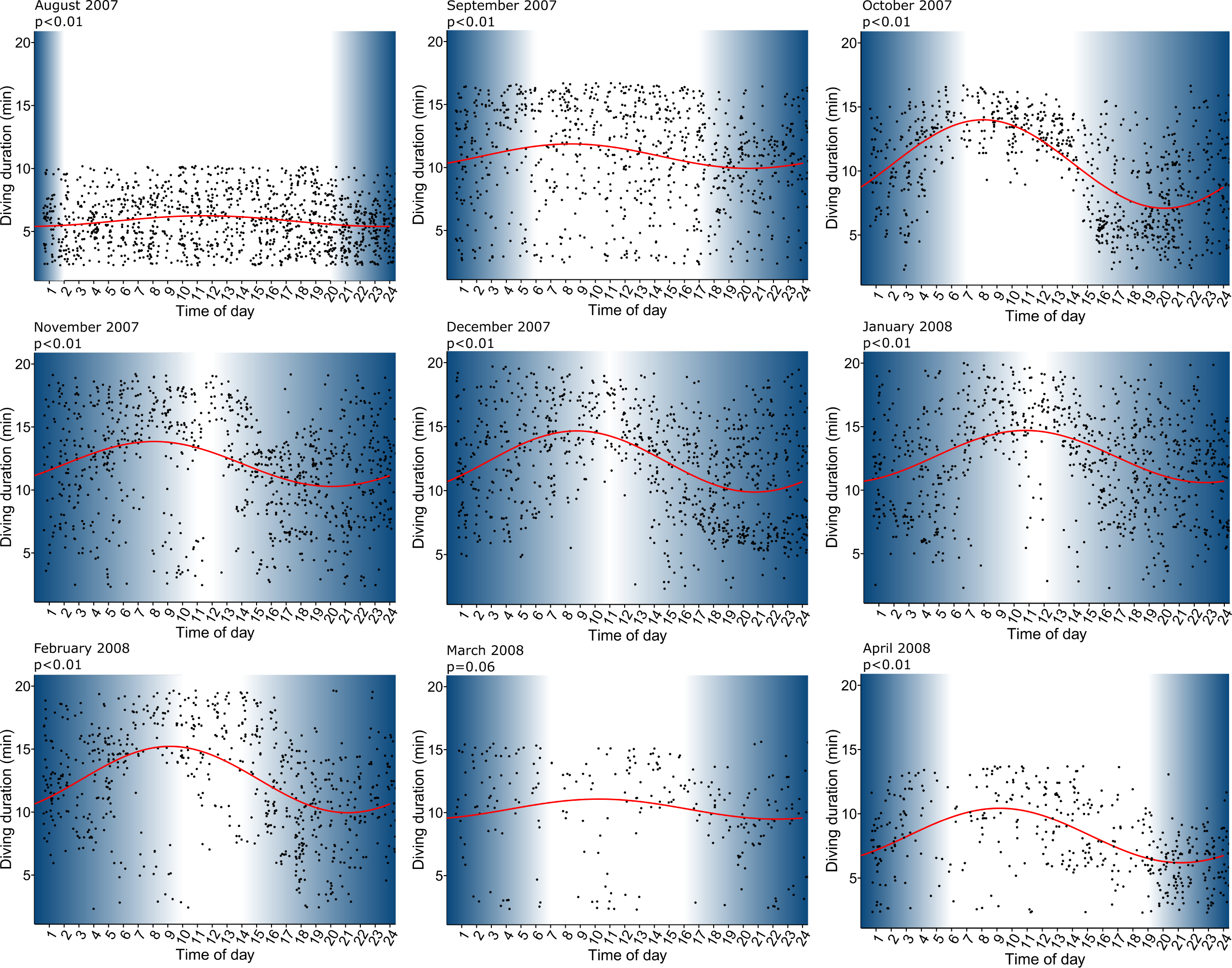
Hourly and daily variations in diving durations in an adult male hooded seal (ID: M171) from the Northeast Atlantic population. Data were collected between August 2007 and April 2008 (data provided by M. Biuw: <u>10.21334/npolar.2017.881dbd20</u>). Only dives lasting between 2 min and the 95 percentiles of maximal duration are included. Each point represents a diving event, the red line represents the cosine curve generated through *cosinor.lm* in RStudio. Each panel corresponds to a different month and the respective p values are indicated in the upper left. Blue and white areas represent day and night hours calculated in relation to position (latitude and longitude) and time of year.

**Figure 2.**
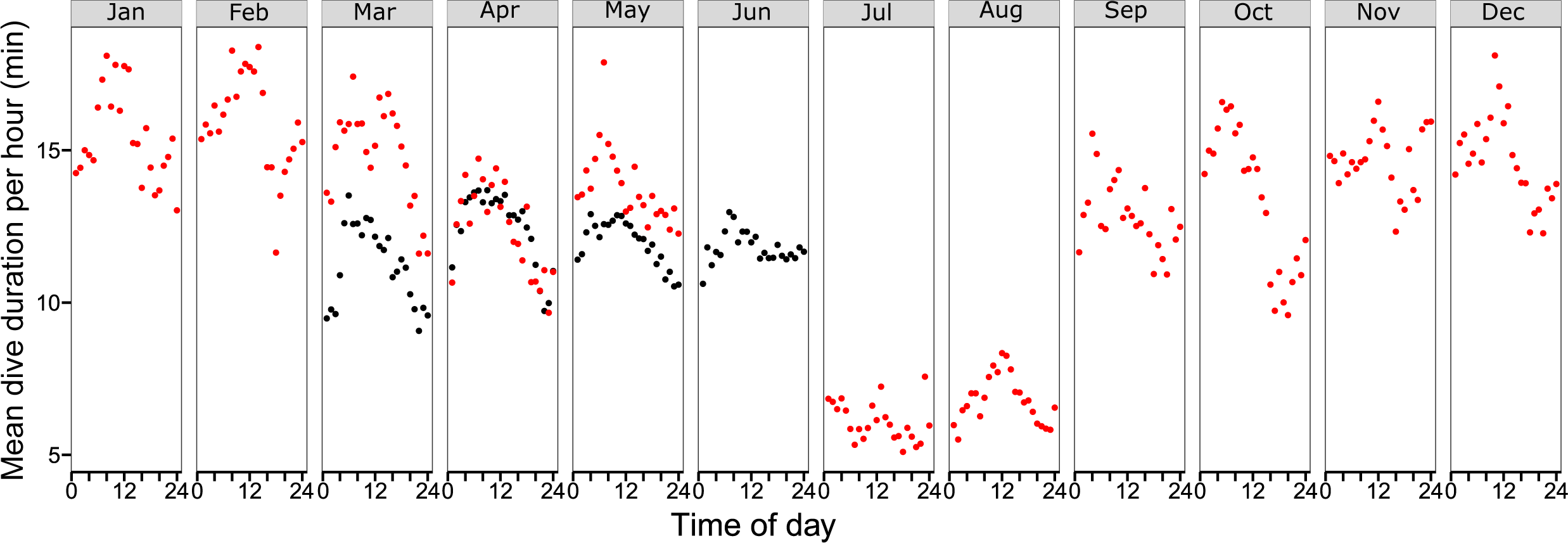
Hourly mean dive duration (minutes) throughout the year for 12 adult hooded seals. Data from 2007 are presented in red (2 adult males); data from 2008 in black (9 adult females and 1 adult male). Only dives lasting between 2 min and 95 percentiles of maximal duration were included in the analysis. Data were binned by hour and the hourly mean dive duration was calculated.

**Table 3.**
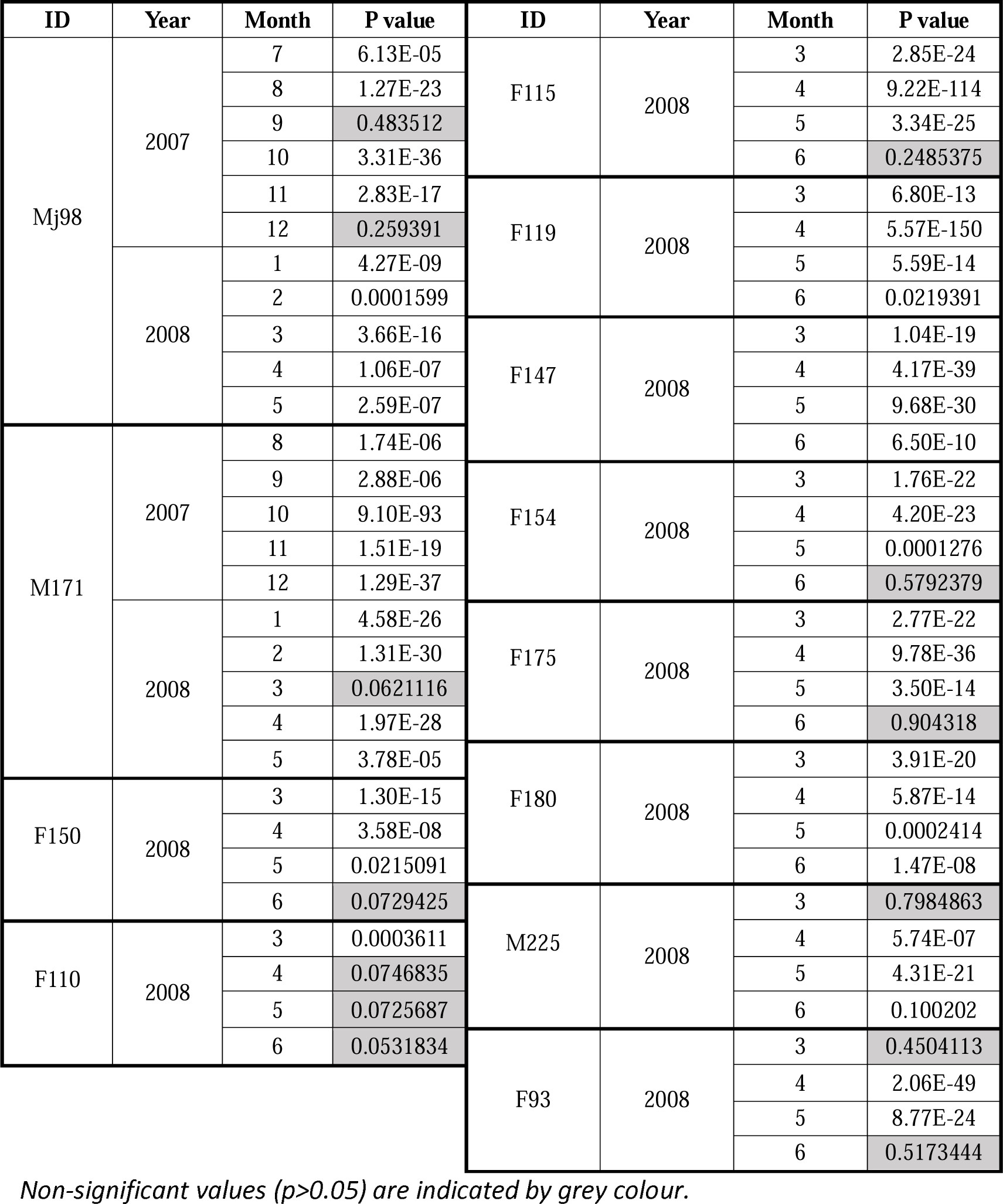
P values for the fitted cosine model for each seal and month calculated through cosinor.detected function in the cosinor2 package in RStudio.

Despite the limited amount of data, we also wished to investigate seasonal changes. We observed that, in accordance to previous data (Folkow and Blix, 1999), the mean value based on the daily distribution of values (MESOR) shows that dives were longer during the winter than in the summer (Fig.S6). Specifically, hourly mean dive duration was particularly short in July and August (Fig. 2) but increased again in September. In October, there was a clear pattern of shorter dives at evening and night and longer dives during the day (Fig. 2), which was persistent throughout all the winter months, until February. In March, dive duration decreased and dives were also less frequent, most likely linked to behavioural changes in connection with the breeding season (Kovacs et al., 1996).

### Circadian molecular clock function in hooded seal fibroblasts and tissues

We derived primary skin fibroblasts from hooded seals and cultured them under a 24-h temperature cycle, to synchronise the cells (Buhr et al., 2010). Measuring the endogenous mRNA abundance of the clock genes *arntl* and *per2*, we show a significant oscillation with a period of 24 hours (JTK cycle, p<0.0001) and antiphase rhythms to one another: *arntl* expression peaking in middle of the low temperature phase and *per2* expression peaking in the middle of the high temperature phase (Fig. 3A). To determine whether these oscillations persist in constant conditions, we synchronised the cells using a temperature cycle and then held temperature constant for 36 hours. Both *arntl* and *per2* endogenous mRNA levels continued to oscillate in anti-phase (Fig. 3B).

**Figure 3.**
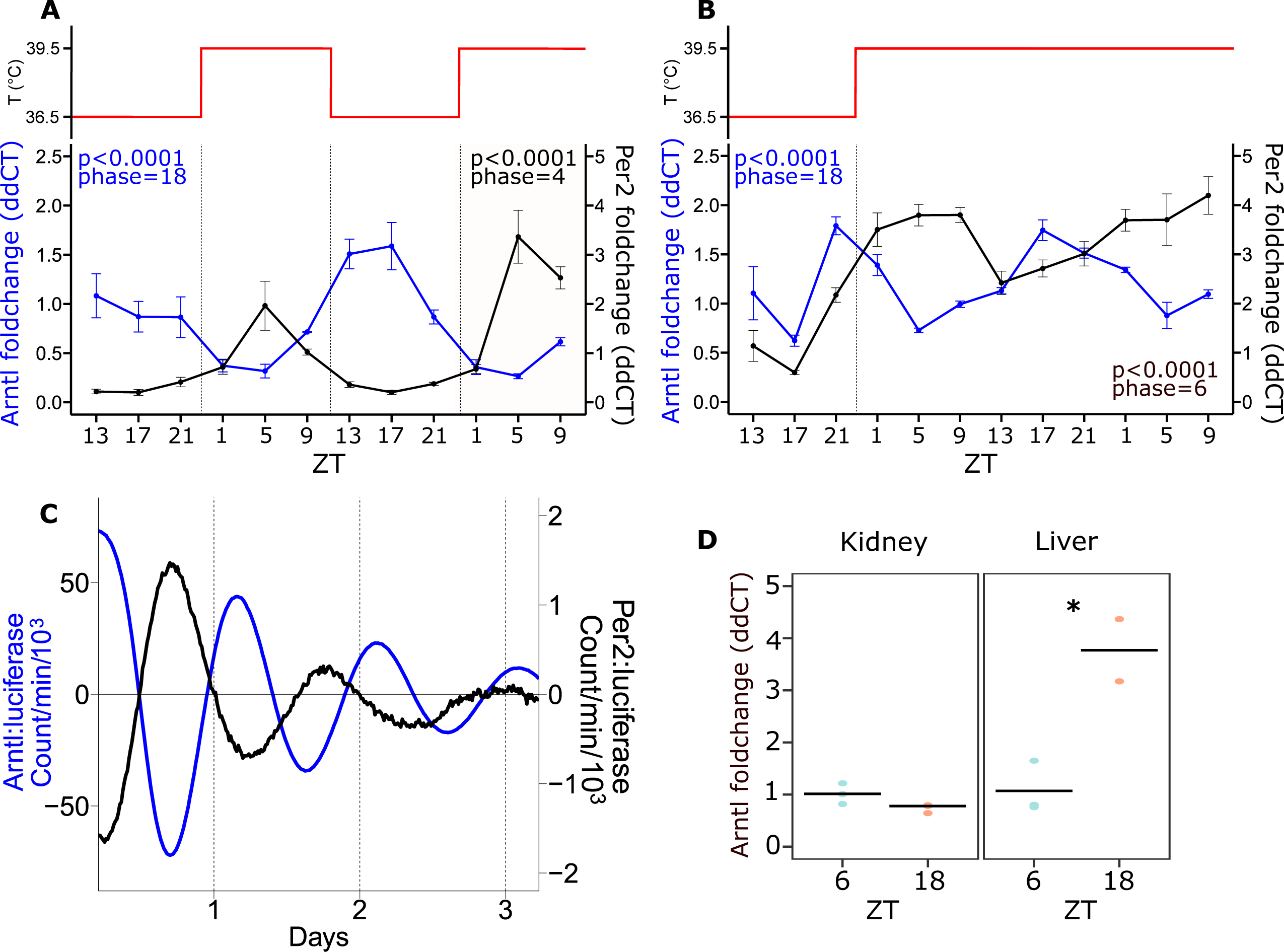
Clock genes expression profile in synchronised hooded seals fibroblasts and tissues. **A)** mRNA expression of core clock genes *arntl* and *per2* in hooded seal fibroblasts synchronised by temperature cycle (T°C), as represented in the upper panel. *Arntl* and *per2* mRNAs show a 24-h period and are in antiphase. P values and phases were calculated through JTK cycle analysis (*n*=4). **B)** mRNA expression of core clock genes *arntl* and *per2* in hooded seal fibroblasts synchronised by temperature cycling and then left at constant temperature (upper panel). *Arntl* and *per2* mRNAs maintain a 24-h period and their antiphasic relationship also in constant conditions. P values and phases were calculated through JTK cycle analysis (*n*=4). **C)** Photon multiplier tube recordings of hooded seal skin fibroblasts transfected with *arntl*:luciferase and *per2*:luciferase. Cells were synchronised with dexamethasone. **D)** *Arntl* mRNA expression in hooded seal kidney and liver tissue, sampled in mid-light phase (ZT6) and mid-dark phase (ZT18). *p<0.05 (One-way ANOVA analysis; *n*=3 except liver ZT18 where *n*=2). Data are expressed as mean ± SEM.

We then generated lentiviral reporters for *arntl* and *per2* to stably transduce our primary fibroblasts and allow real time monitoring of clock gene expression for multiple days. After synchronisation with dexamethasone, we observed significant oscillations of approximately 24 hours for both *arntl* and *per2* (R^2^ = 0.96 and 0.95, respectively) and persistent oscillations for 3 days post synchronisation (Fig. 3C). These results are consistent with a functional molecular clockwork in hooded seal fibroblasts.

We also measured endogenous *arntl* mRNA expression in liver and kidney samples collected in the mid-light (ZT6) and mid-dark phase (ZT18) from captive hooded seals held on a LD12:12 light-dark cycle. *Arntl* expression in the liver showed a significant time-of-day effect, with expression at ZT18 being some four-fold higher than at ZT6 (p<0.05; Fig. 3D).

### Circadian clock phase relationship with the metabolic capacity of hooded seal mitochondria

To investigate whether there is a time-of-day variation in mitochondrial respiration, we used a O2k high-resolution respirometers (Oroboros) substrate-uncoupler-inhibitor titration (SUIT) protocol (see Table 2) with permeabilised skin fibroblasts from the hooded seal. We sampled at 6 different zeitgeber times (ZT) across a 48-h temperature cycling experiment (Fig. S1A). We used PCA analysis to integrate values for the 7 parameters measured by the respirometers for each individual sample taken at each time-point. This generated 2 principal components (PC1 and PC2), which together accounted for over 90% of the overall variance for these 7 parameters across the study as a whole (Fig. 4A). There was wide ZT-dependent variation, and we speculated that this variation was related to predicted phases for *arntl* and *per2* expression. Based on the endogenous mRNA expression (Fig. 3A) we focused on ZT5 (low *arntl*, high *per2* expression) and ZT17 (high *arntl*, low *per2* expression), which separated more clearly on the PCA plot (Fig. 4A). Among all the mitochondrial states, the *leak state* at complex I (LeakN; Table 2) was the only one to show a statistically significant difference between ZT5 and ZT17 (p=0.001). We therefore calculated complex I coupling efficiency using Eq. 1 across all the ZTs.

**Figure 4.**
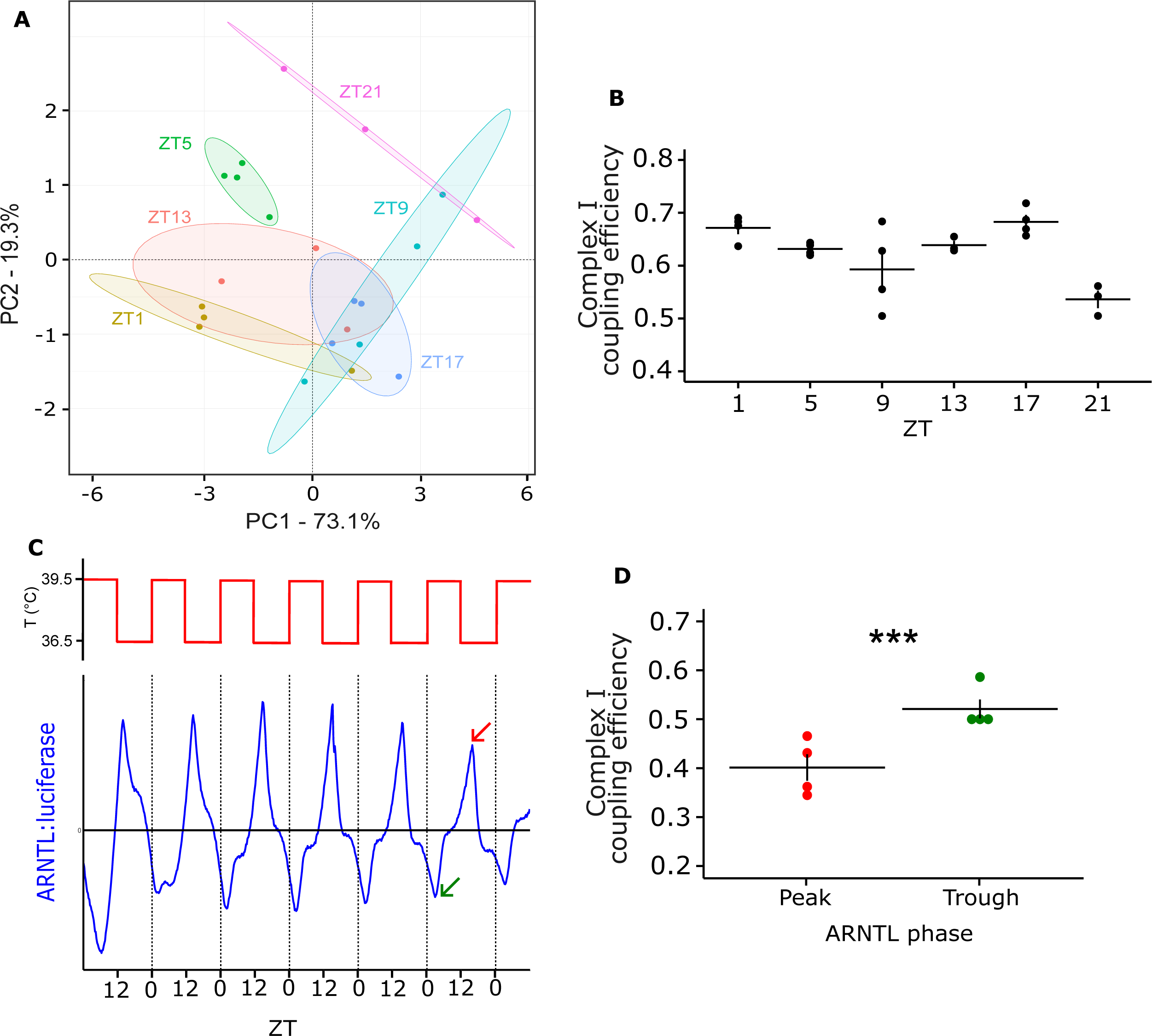
Relationship between clock gene expression rhythms and mitochondrial respiration in hooded seal fibroblast cultures. **A)** PCA analysis across 7 mitochondrial states (*Leak* (LeakN, LeakOmy), *OXPHOS* (PM-P, CI-P, CI +CII-P, CII-P), *Uncoupled state* (CII-E)). There is strong clustering at ZT5 and ZT17. Ellipses were drawn with a 0.80 confidence interval (*n*=4 except at ZT13, 21 where *n*=3). **B)** Complex I coupling efficiency (calculated as 1-(LeakN/PM-P)) at all the ZTs where oxygen consumption measurements were made. Coupling efficiency shows a time-dependent variation (one-way ANOVA: p<0.01; Kruskal-Wallis test: p<0.05; *n*=4 except ZT13, 21 where *n*=3) **C)** Photon multiplier tube recordings of hooded seal skin fibroblasts transfected with *arntl*:luciferase and synchronised by temperature cycling (36.5 – 39.5°C), as indicated at the top. Recordings from the first 5 days were used to calculate expected peak (red arrow) and through (green arrow). **D)** Complex I coupling efficiency calculated as in **B,** at the expected peak and through indicated in **C** (One way ANOVA: p<0.001; Mann-Whitney U-test: p<0.05). Data are expressed as mean ± SEM. ***p<0.001 (One-way ANOVA analysis, *n*=4).

This revealed the presence of a time effect on complex I efficiency (p<0.01, One-way ANOVA; p<0.05, Kruskal-Wallis test; fig. 4B), with lowest values at ZT21 (0.46 ± 0.017). High coupling efficiencies (close to 1) indicate highly coupled mitochondria (Doerrier et al., 2018; Gnaiger, 2020) and changes in coupling efficiency at complex I suggest a time of day effect on mitochondrial function.

To provide independent verification of the relationship between mitochondrial activity and clock phase we collected samples from virally transduced *arntl:*luciferase-reporter-expressing hooded seal fibroblasts at phases of peak and trough *arntl* expression and processed them for respirometry analysis (Fig. 4C). Calculation of complex I coupling efficiency revealed a lower oxidative phosphorylation activity at complex I at peak *arntl* expression (0.40 ± 0.024) than at trough (0.52 ± 0.017)(p<0.001, One-way ANOVA; p<0.05 Mann-Whitney U-test; Fig. 4D). Therefore, when *arntl* is at its peak, complex I is not as efficiently coupled as when *arntl* is at its trough, indicating a clock-dependent variation of complex I coupling efficiency. These difference in coupling efficiency may relate to regulation of ROS production or oxidative metabolism.

## DISCUSSION

The present study confirms that under natural conditions hooded seal diving behaviour is highly time of day dependent, and further demonstrates that under laboratory conditions, hooded seal cells show robust circadian characteristics, which are associated with cyclical changes in mitochondrial activity. Daily physiological and molecular changes are a key factor for determining the ability of different species to overcome the challenges imposed by a continuously changing environment. For the hooded seal the biggest challenge probably is the restricted access to oxygen during long and deep dives.

We report that dives were consistently longer during the day and shorter at night and that there was a 24-h period in the diving duration pattern. Such daily variations are most likely related to similar patterns in the behaviour of hooded seal prey (e.g., by redfish (*Sebastes spp.)* and squid (*Gonatus spp.*)), which can display diurnal vertical migrations as their prey, in turn, migrate according to the photic state in the water column (Kristensen, 1984; Torsvik et al., 1995). Even though the day-night trend is quite consistent, there is some seasonal variation, as previously reported (Andersen et al., 2013). These variations relate in part to the breeding season (late March) and the moulting season (June/July), during which adult hooded seals spend more time hauled out (Folkow and Blix, 1999; Folkow et al., 1996; Kovacs et al., 1996; Øritsland, 1959). However, dives also tend to generally be deeper and longer in winter than in summer (Folkow & Blix, 1999). While the ultimate causation for diel patterning (i.e. prey availability) is clear, the proximate mechanisms dictating the time-of-day dependent diving behaviour is unknown, this led us to investigate whether a circadian clock – mitochondria coupling might be present in hooded seal cells.

Here, we show that circadian changes in *arntl* expression in hooded seal fibroblasts coincide with a significant change in the coupling efficiency of complex I. When *arntl* expression is at its trough, complex I coupling efficiency is high. In support of this view we show, by two independent methods, that primary hooded seal skin fibroblasts display antiphase circadian oscillations of the clock genes *arntl* and *per2*, a result consistent with other characterisations of the mammalian circadian clock (Buhr and Takahashi, 2013). These oscillations are circadian because they persist under constant conditions (temperature, in this study) and with a period of approximately 24 hours (Dunlap et al., 2004). We also show that the activity of complex I varies according to time-of-day and to the circadian clock phase defined by *arntl*.

Diurnal oscillations in complex I activity have been documented in mice liver (Jacobi et al., 2015; Neufeld-Cohen et al., 2016) and in human skeletal muscle (van Moorsel et al., 2016). In the human cell line HepG2, *arntl* expression regulates complex I activity by a process of acetylation and deacetylation through the NAMPT-NAD-SIRT1/3 machinery (Cela et al., 2016).

Finally, we show a time-of-day difference in *arntl* gene expression in the hooded seal liver, consistent with previous recordings in freshly isolated mice livers (Jacobi et al., 2015; Manella et al., 2020). However, we observed no difference in the kidneys, but we were limited to only two time points so no strong conclusions regarding tissue-specific circadian rhythmicity can be drawn. We wanted to consider those organs specifically, because they are known to undergo substantial reduction in their arterial blood supply (by >80%) – and, hence, in the supply of blood-borne O_2_ - in connection with long-duration diving in seals (e.g. Blix et al., 1983; Zapol et al., 1979). Therefore, the liver and kidney appear to be particularly hypoxia-/ischemia-tolerant (e.g., Hochachka et al., 1988), from which follows that a potentially tissue-dependent circadian clock phasing and interaction with mitochondria may be expected to be found in these particular tissues. However, further studies are needed to investigate this.

In summary, we have identified a circadian clock phase-dependent change in complex I coupling efficiency, demonstrating for the first time a mitochondria – clock interaction in the hooded seal. This change in coupling efficiency is determined by clock-dependent changes in complex I leak state and may represent a switch from an OXPHOS-based metabolism to less oxidative metabolism. But it could also be interpreted as a protective mechanism, to modulate the amount of reactive oxygen species (ROS) produced by the mitochondria through regulation of complex I leak (Brand, 2000; Cadenas, 2018). We speculate that the existence of mitochondria - circadian clock coupling that regulates either ROS production or oxidative metabolism *may* enhance the ability of seals to tolerate long-duration dives, by providing additional hypoxia tolerance mechanisms at the time it is needed the most - i.e., during the day, when they forage at greater depths and for longer durations (Fig. 1). However, a functional link remains to be determined. Our findings suggest that future experiments should take circadian clock phase into account, when investigating mitochondrial responses to hypoxia and the role of HIF-1 in diving mammals.

## ACKNOWLEDGMENTS

The authors would like to thank the animal technicians of the Arctic Chronobiology and Physiology group for all the help with the animal handling: Hans Lian, Hans-Arne Solvang, and Renate Thorvaldsen. We would also like to thank Dr. Martin Biuw for his help in the initial stages of the dive data analyses and Dr. Daniel Appenroth for his work on the lentiviral reporter.

## AUTHOR CONTRIBUTIONS

Conceptualization: S.H.W., A.C.W, D.G.H., C.C.; Methodology: A.C.W., F.K., C.C.; Software: C.C.; Validation: F.K., C.C.; Formal analysis: C.C., D.G.H., A.C.W.; Investigation: C.C., F.K., A.C.W., S.H.W.; Resources: S.H.W., L.P.F.; Data curation: C.C.; Writing-original draft: C.C., D.G.H.; Writing – review and editing: D.G.H., C.C., S.H.W., A.C.W., L.P.F., F.K.; Visualization: C.C., D.G.H., A.C.W.; Supervision: L.P. F., S.H.W., A.C.W., D.G.H.; Project administration: S.H.W., L.P.F; Funding acquisition: S.H.W.

## DATA AVAILABILITY

Code used for the analysis and data are available at https://github.com/ShonaWood/SealClock

