## Supplementary figures for "Circadian coupling of mitochondria in a deep-diving mammal"

**Supplemental figures**

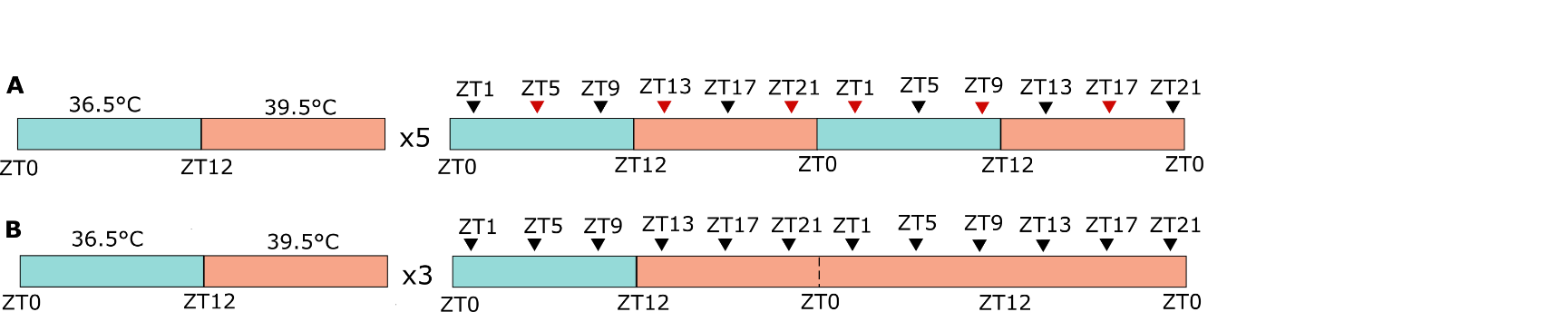

**Figure S1. Experimental design of temperature treatments used on hooded seal skin fibroblasts.** **A)** Temperature cycling treatment: cells were exposed to 5 temperature cycles before being collected every 4 hours. Sampling start is indicated by arrow heads. Black arrow heads represent RNA sampling, red arrow heads represent RNA and mitochondrial oxygen consumption sampling. **B)** Constant temperature treatment: cells were exposed to 3 cycles of cycling temperature and then sampled at constant temperature. Only samples for RNA extraction were taken in this experiment. ZT= Zeitgeber Time.

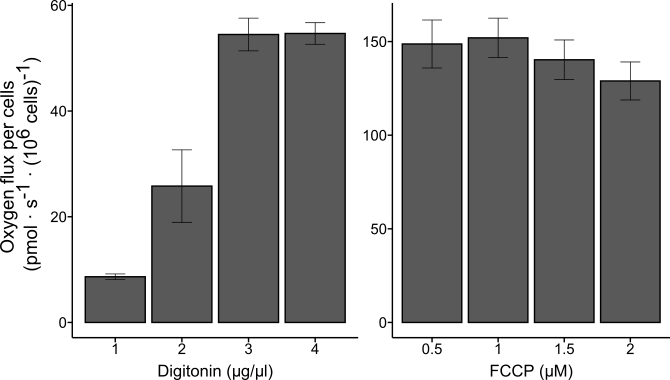

**Figure S2. Optimal concentration of digitonin and FCCP.** Prior to the main experiments, the optimal concentrations of digitonin and FCCP were determined in separate pilot experiments. For digitonin, the SUIT-010 (Doerrier et al., 2018) was followed: optimal concentration was assumed when maximal O_2_ flux was recorded in the chamber. For FCCP, cells were treated with multiple FCCP titrations until O_2_ flux started to decrease. Data are represented as mean ± SEM and expressed as picomoles of O_2_ per second per million cell (10^6^ cells).

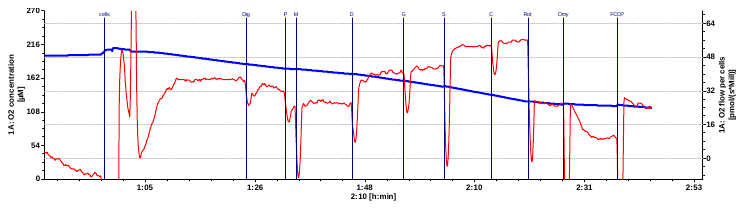

**Figure S3. Original oxygraph from mitochondrial respiration measurements.** The oxygraph shows the real time changes in O_2_ flow per cell (pmol ^●^  s^-1^ ^●^  (10^6^ cells)^-1^) in red and the O_2_ concentration in the Oroboros oxygraphic chamber (µM) in blue. Vertical blue lines represent injections following the SUIT protocol explained in Table 2. Cells: insertion of cell sample in the oxygraphic chamber; Dig: digitonin; P: pyruvate; M: malate; D: ADP; G: glutamate; S: succinate; C: cytochrome c; Rot: rotenone; Omy: oligomycin; FCCP: uncoupler.

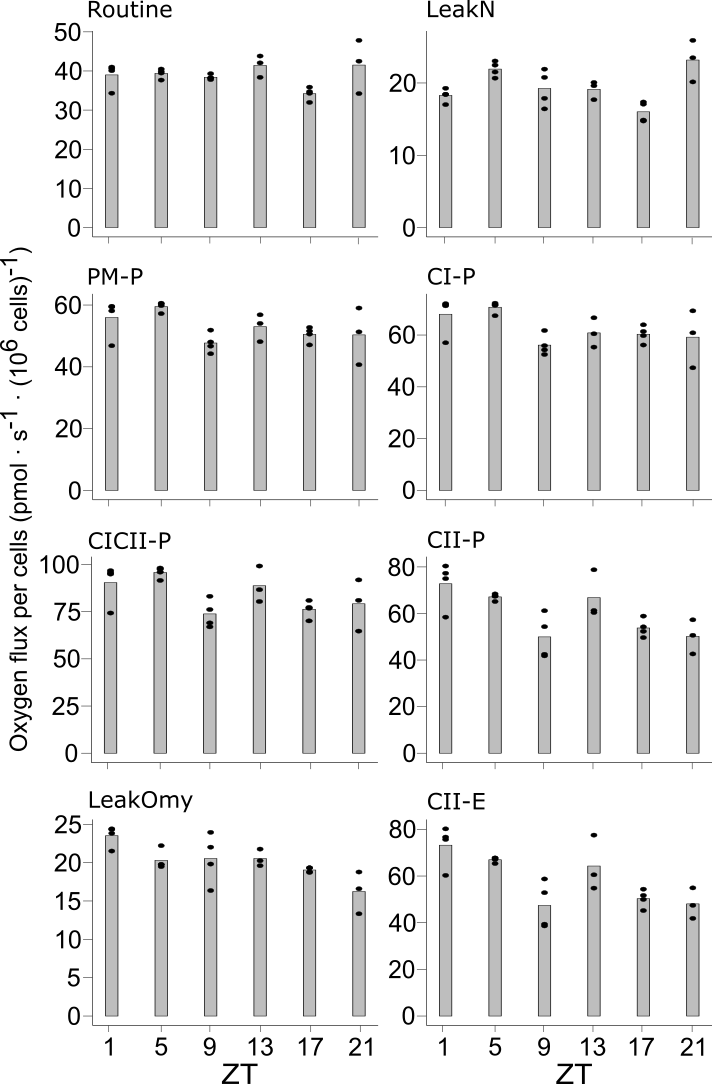

**Figure S4. Average oxygen flux per each mitochondrial state**. The average measures for each mitochondrial state are shown in the figure as picomoles (pmol) of O_2_ per second per million cell (10^6^). Routine: basal respiration before addition of any substrate; LeakN: leak state at complex I in the presence of pyruvate and malate; PM-P: OXPHOS through complex I in the presence of pyruvate, malate and ADP; CI-P: OXPHOS through complex I in the presence of pyruvate, malate, glutamate and ADP; CICII-P: OXPHOS through complex I and complex II in the presence of pyruvate, malate, glutamate, succinate and ADP; CII-P: OXPHOS through complex II after inhibition of complex with rotenone I; LeakOmy: leak state at complex II after inhibition of ATP synthase with oligomycin; CII-E: uncoupled state measuring maximal mitochondrial capacity after addition of FCCP.

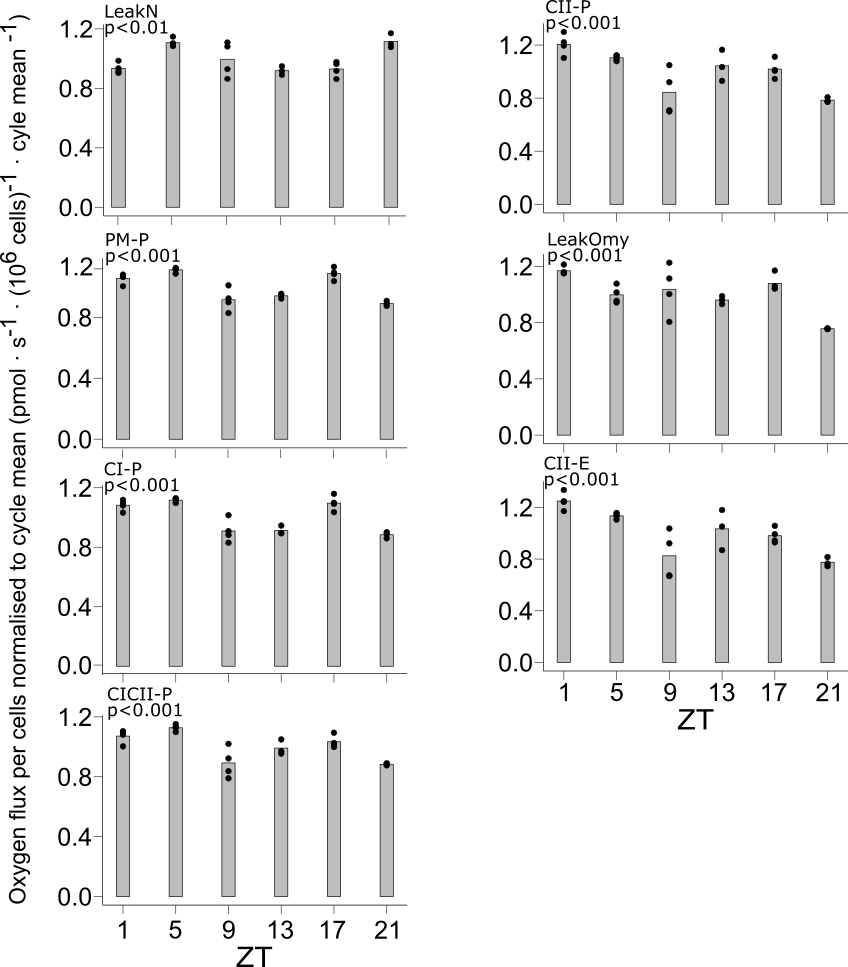

**Figure S5**. **Mitochondrial states normalised to cycle mean.** Respiratory rates were first normalised to million cells (10^6^) in DatLab software. Then, they were normalised to ‘Routine respiration’, defined as basal respiration without addition of any substrate (Table 2). Finally, the cycle mean was calculated and used to normalise the respiratory rate, expressed as a ratio of the mean, and represented as oscillations around the value 1. P-values are indicated as calculated across all the ZTs with one-way ANOVA. (Total *n*=22, with *n*=4 for each ZT except at ZT13, 21 where *n*=3). LeakN: leak state at complex I in the presence of pyruvate and malate; PM-P: OXPHOS through complex I in the presence of pyruvate, malate and ADP; CI-P: OXPHOS through complex I in the presence of pyruvate, malate, glutamate and ADP; CICII-P: OXPHOS through complex I and complex II in the presence of pyruvate, malate, glutamate, succinate and ADP; CII-P: OXPHOS through complex II after inhibition of complex with rotenone I; LeakOmy: leak state at complex II after inhibition of ATP synthase with oligomycin; CII-E: uncoupled state measuring maximal mitochondrial capacity after addition of FCCP.

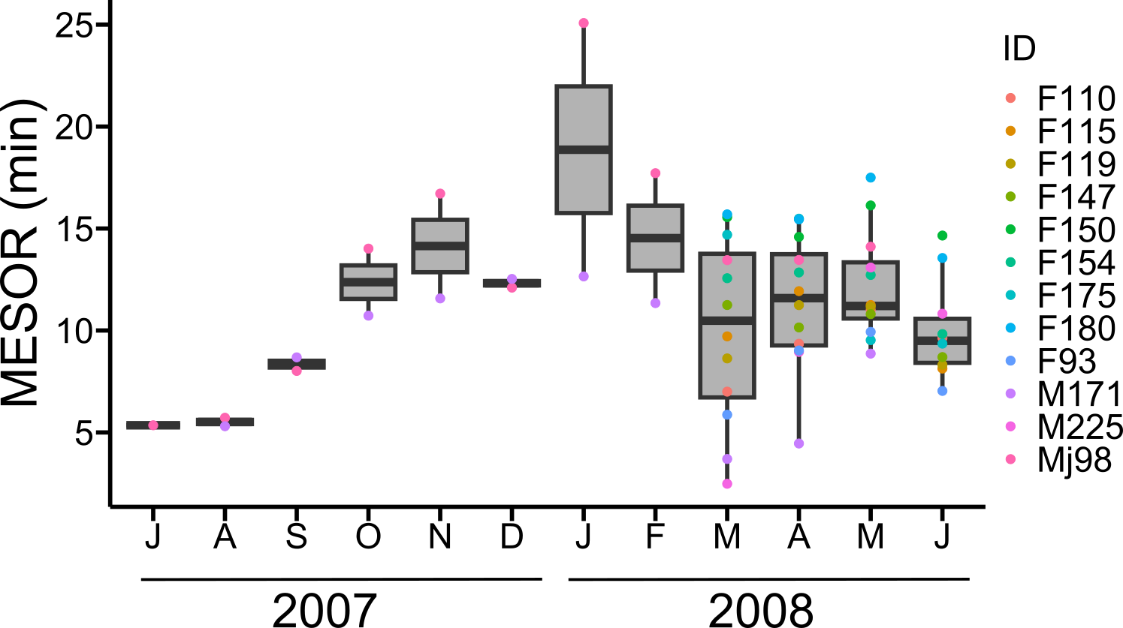

**Figure S6. Dive duration MESOR calculated for each seal and for each month from July 2007 until June 2008**. In the legend, the seals IDs are listed with the gender (F=female, M=male) and their body weight. Only dives lasting between 2 min and 95 percentiles of maximal duration were considered in the analysis.
